# Express detection and discrimination of visual objects by primate superior colliculus neurons

**DOI:** 10.1101/2022.02.08.479583

**Authors:** Amarender R. Bogadhi, Ziad M. Hafed

## Abstract

Primate superior colliculus (SC) neurons exhibit rich visual feature tuning properties and are implicated in a subcortical network hypothesized to mediate fast threat and/or conspecific detection. However, the mechanisms through which generalized object detection may be mediated by SC neurons remain unclear. Here we explored whether, and how quickly, SC neurons detect and discriminate real-life object stimuli. We presented experimentally-controlled gray-scale images of seven different object categories within the response fields (RF’s) of SC neurons, and we also presented a variety of luminance- and spectral-matched image controls. We found that all of our functionally-identified SC neuron types preferentially detected real-life objects even in their very first stimulus-evoked visual bursts (starting within approximately 50 ms from image onset). Intriguingly, even visually-responsive motor-related neurons exhibited such robust early object detection, and they also preferentially discriminated between object categories in their initial visual bursts. We further identified spatial frequency information in visual images as a critical source for early object detection and discrimination by SC neurons. Our results demonstrate rapid and robust SC processing of visual objects, and they underline how the visual processing capabilities of the primate SC support perception and action.

## Introduction

Object detection and recognition are fundamental components of primate vision, and a substantial number of visual cortical areas are dedicated to processing visual objects [1-5]. However, vision does not occur in complete isolation of behavior, and an element of visual object processing in the brain must facilitate active orienting in association with objects, whether to avoid threats [6] or to foveate and further process behaviorally-relevant items. Indeed, certain classes of visual objects, like faces, easily pop out from visual scenes with very short latencies [7], and short-latency eye movements can likewise be automatically captured by completely task-irrelevant object images [8].

The speed with which orienting phenomena associated with visual object recognition proceed points to the presence of subcortical mechanisms for visual object processing. Indeed, in 1974, Updyke [9] observed neurons in the superior colliculus (SC), a site of convergence for retinal and extra-retinal visual signals, that were particularly sensitive to three-dimensional objects, and SC cells sensitive to complex visual stimuli were also reported by Rizzolatti and colleagues in 1980 [10]. More recently, a series of studies explored the roles of the SC and pulvinar in the processing of face and snake images [11-17], and concluded that the SC may be part of a fast detection network for visual threats and ecologically-relevant faces that can influence emotions [6].

Because the SC is also shown to contribute to a variety of important cognitive processes like target selection, visual attention, and perceptual decision making [18-24], and given that SC activity can influence cortical areas through different thalamic circuits [25-28], it stands to reason that the SC may be involved in object processing in a more general way than being specifically tuned for processing snakes and faces. In fact, experimental manipulation of SC activity is associated with altered object selectivity in a patch of the ventral visual processing stream of the cortex [29], and, similarly, the SC has a dedicated primary cortical area in mice [30]. Importantly, the SC possesses privileged access to the saccadic system’s motor periphery; therefore, a generalized object detection capability by the SC can support rapid orienting behaviors, which are facilitated by visual objects [8]. As a result, there is a pressing need to investigate whether, and how, neurons in the primate SC detect and discriminate visual objects. We did so by presenting seven different categories of visual object images to individual SC neurons, along with various versions of control images. We found rapid and sustained detection and discrimination of visual object categories by all visually-responsive SC neuron types. We also observed that SC tuning to spatial frequency information in images [31, 32] can facilitate the fastest components of SC visual object processing. Generalized visual object processing is a robust property of the primate SC.

## Results

### The very first SC visual responses differentiate between object and non-object stimuli

We analyzed SC visual responses to images of real-life objects appearing within the recorded neurons’ response fields (RF’s). The monkeys fixated a central spot, and we presented one of 28 different images, drawn from seven different object categories and their corresponding control images (Figs. 1A, S1). The control images were luminance- and spectrum-matched non-object images (Methods): phase-scrambled controls had the same spatial frequency content as the real object images, but with spectral phase scrambling; grid-scrambled images had small, square patches (grids) containing identical copies of small patches from the original images, but with randomized locations. The grid-scrambled images maintained local image properties but disrupted global form information. Finally, since grid scrambling also introduced a square grid of hard edges between the scrambled image patches (altering the spatial frequency content of the images), we also checked whether object detection by SC neurons was significantly disrupted by overlaid grids presented over the intact objects (grid+object images; Figs. 1A, S1). Thus, each neuron was tested with seven different object categories and four different image types: two being coherent objects (object and object+grid) and two being image-matched, non-object images (grid-scrambled and phase-scrambled).

**Figure 1.**
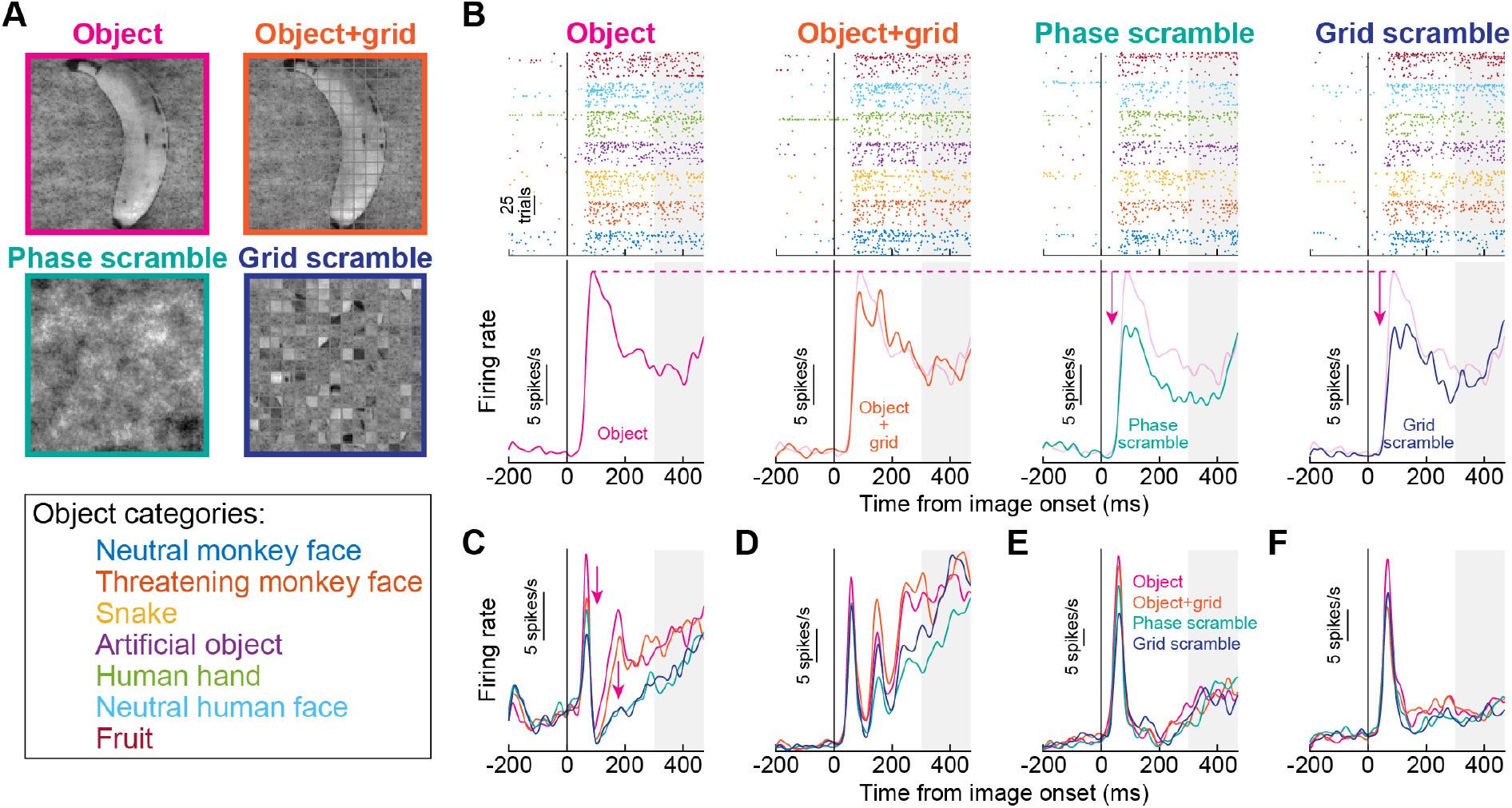
Early and sustained enhancement of superior colliculus (SC) visual activity for real-life object images. **(A)** We presented an image of real-life objects (e.g. banana) as well as multiple variants of it. The object+grid image overlaid a grid creating small square patches of image regions. The phase-scrambled image contained the same spatial frequency content as the object image, but with scrambled phase information. And, the grid-scrambled image had randomized grid locations from the object+grid image. In total, we tested seven different object categories, spanning faces, animals, and artificial objects. See also Fig. S1. **(B)** Each column shows the responses of an example neuron under the four different image conditions. The leftmost column shows responses to intact object images. Top: individual trial spike time rasters showing responses to each object category; bottom: average firing rate plot pooling the seven different object categories together (but see Figs. 4, 5 later for further analyses of object preference without pooling). The neuron exhibited a robust visual burst followed by sustained activity. In the second column, the overlaid grid minimally altered the response. However, both the phase-scrambled (third column) and grid-scrambled (fourth column) conditions were associated with significantly weaker activity. **(C-F)** Four additional example neurons showing similar results. The object and object+grid conditions had the highest initial visual bursts. Moreover, sustained activity was higher for the object and object+grid conditions than for the scrambled conditions. The gray shaded regions in **B**-**F** denote the time at which the saccade target could appear in the subsequent stages of the trials (Fig. S1).

Initial and sustained SC visual responses were systematically higher for real object stimuli than for non-object images. Consider the example neuron of Fig. 1B. In both the object and object+grid conditions (leftmost two columns), the neuron’s visual response was higher than in the phase- and grid-scrambled conditions (rightmost two columns). Therefore, the neuron discriminated between intact object and non-object stimuli even within its very first, initial visual burst (i.e. within approximately 50 ms from image onset).

In Fig. 1C-F, we also show results from four additional example neurons. In all cases, initial visual bursts were the highest for real object images and/or object+grid images. Moreover, sustained visual activity was clearly higher for the object and object+grid images than for the phase- and grid-scrambled images, and this was the case even for the neurons with relatively low sustained activity (Fig. 1E, F). Note that in these analyses, we pooled all seven object categories together, but we later return to the question of whether SC neurons also preferred specific individual objects or not. Also note that starting at 300 ms after image onset (gray shaded regions), the saccade target could appear for the next stages of the behavioral task (Methods). Therefore, in all subsequent analyses, we focused only on the first 300 ms of neural responses. In all, the five example neurons of Fig. 1 suggest the presence of both very early as well as sustained discrimination of object and non-object stimuli by primate SC neurons.

We confirmed that, across the population, early SC visual bursts robustly discriminated between object and non-object images. We did so by assessing the discriminability of firing rates between the object and grid-scrambled conditions; we performed a running receiver operating characteristic (ROC) analysis on the neural responses, using 40 ms time bins in steps of 10 ms (Methods). For each time bin around image onset, we collected firing rates from each condition (either intact object or grid-scrambled image) pooled for all seven object categories, and we then calculated the area under the ROC curve (AUC) between the two distributions (see later for our separate analyses of object preference). AUC values significantly different from 0.5 indicated discriminable firing rate distributions between object and grid-scrambled images (Methods).

In each monkey, we accepted a neuron as significantly detecting objects versus non-object stimuli if it had a significant AUC value in any time bin within 0-300 ms from image onset (Methods). Out of 131 neurons in monkey M (including task-irrelevant ones like purely motor neurons), 77 showed significant discrimination performance for intact objects relative to grid-scrambled images. In monkey A, 26 out of 82 total neurons (again including task-irrelevant ones like purely motor neurons) did so. Most importantly, in both monkeys, the highest discrimination performance always occurred in the very initial visual burst interval. This is illustrated in Fig. 2A for monkey M and Fig. 2D for monkey A (error bars denote 95% confidence intervals). Therefore, SC neurons detect visual objects in an express manner, consistent with behavioral evidence of an automatic influence of visual forms on target selection for eye movements [8], and also consistent with results demonstrating altered cortical object selectivity with altered SC activity [29].

**Figure 2.**
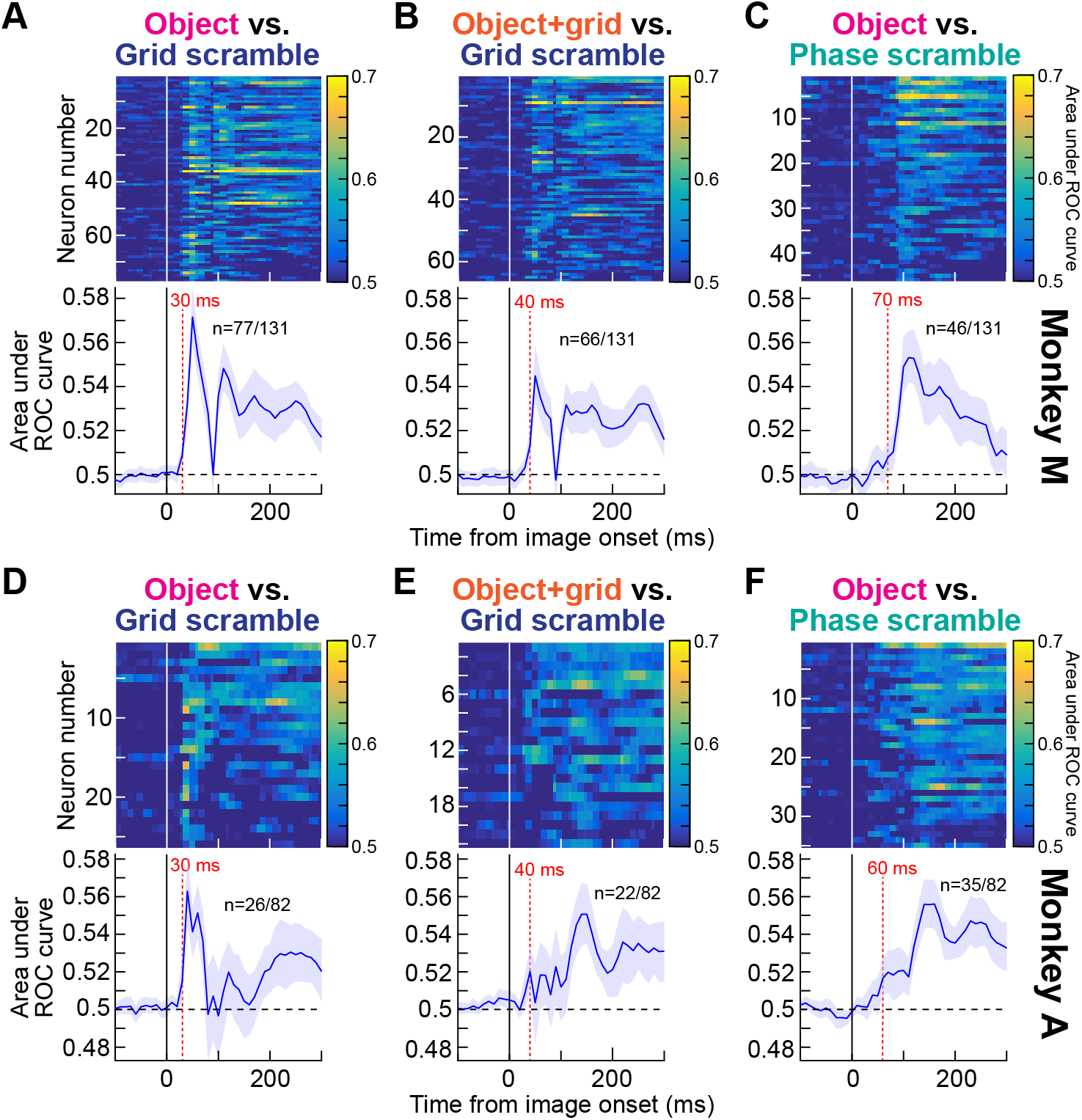
Early and late detection of visual objects by SC neurons. **(A)** For each neuron in monkey M, we compared distributions of firing rates (in 40 ms time bins) between intact and grid-scrambled object images using ROC analyses (Methods). For each neuron with a significant AUC (area under ROC curve) value in the interval 0-300 ms from image onset (n=77), we plotted AUC as a function of time in the top panel (the color indicates the AUC value). The bottom panel plots the average of all neurons’ AUC time courses (error bars denote 95% confidence intervals across the population), showing an initial robust peak followed by sustained elevation. The dashed vertical line marks the first time point after stimulus onset for which the AUC value of the population was significantly deviated away from 0.5 (30 ms). **(B)** Same analysis but comparing object+grid images to grid-scrambled images. The overlay of a grid on top of the images (Fig. 1A, S1B) was not enough to strongly alter the ability of the neurons to detect visual objects, but the altered spatial frequency content of object+grid images slightly modified the early (<100 ms) AUC values (see **C**). **(C)** Same analysis but comparing object images to phase-scrambled images. Here, the early peak in AUC discrimination performance was significantly attenuated, suggesting that the spatial frequency content of object images contributes to early object detection mechanisms by the SC. **(D-F)** Same as **A**-**C** but for monkey A. The results in both animals were highly consistent with each other. Figure S2 shows related analyses controlling for the effects of microsaccades.

Since grid scrambling necessarily entailed adding hard vertical and horizontal edges around each grid (see the example grid-scrambled image in Fig. 1A), we also checked whether the results of Fig. 2A, D were trivially explained by these added edges. We, therefore, repeated the ROC analyses, but this time comparing the grid+object images to the grid-scrambled ones. Now, both image types had the same hard edges embedded within them, but the grid+object images preserved much of the form information in the original intact object images; the grid+object stimuli were akin to the objects being occluded by a thin rectangular mesh and thus still recognizable as coherent objects. We still found robust early and sustained discrimination performance in both monkeys (Fig. 2B, E). Thus, the results of Fig. 2A, D were not explained by the slightly altered spatial frequency content introduced by the grids in the grid-scrambled images. We next explored spatial frequency effects more closely.

### The earliest phase of visual-object detection by SC neurons relies on spatial frequency image content

Because spatial frequency is relevant for visual object recognition [33-37], and because primate SC neurons exhibit spatial frequency tuning [31, 32], we next asked how object detection performance as in Figs. 1, 2A, 2B, 2D, 2E depended on spatial frequency. We repeated the ROC analyses, but we now pitted intact object images against phase-scrambled images (Methods). In these latter images, there was no grid overlay, but the phases of the different spatial frequency bands of the images were randomized relative to the intact object image condition. We still found a substantial number of neurons in each monkey with significant AUC values in the first 300 ms after image onset (Fig. 2C, F), satisfying our criteria for object detection by SC neurons. Interestingly, however, the earliest phase of AUC discrimination performance between intact and phase-scrambled images was significantly weaker than in the case of grid scrambling. For example, across the population of significant neurons in each monkey in the phase-scrambled condition (46 in monkey M and 35 in monkey A), the average population AUC value first moved significantly away from 0.5 (at the 95% confidence level) at 70 ms and 60 ms after image onset for monkeys M and A, respectively (Fig. 2B, E). This is in contrast to the earlier detection of objects with respect to grid-scrambled images (30 ms; Fig. 2A, D). This observation implies that in the very early phases of visual responses in our population, neural activity for the intact objects was more similar to that for phase-scrambled objects than it was to grid-scrambled images. Therefore, object detection by SC neurons is partially mediated, in the very early phases of neural responses, by the spatial frequency image processing capabilities of these neurons; this highlights an interesting potential functional role for spatial frequency tuning in primate SC neurons [31, 32].

It is, nonetheless, interesting that in longer intervals after image onset (e.g. >100 ms), there was still significant AUC discrimination performance between the intact and phase-scrambled object images. This is clearly seen in Fig. 2C, F, in which significant AUC discrimination performance persisted at least until the next phase of the trials (>300 ms). Such sustained effect might suggest a reverberation of object representation between the SC and other visual cortical areas associated with object recognition. For example, because object recognition may preferentially benefit from mid-spatial-frequency information [34-37] and the SC is primarily low-spatial-frequency tuned [31], feedback to the SC after the initial visual bursts can help to stabilize the SC representation for the detected objects for prolonged intervals. Therefore, object detection by SC neurons proceeds with both an early and a sustained phase (Fig. 2A, D); the early phase is supported by spatial frequency information that is intrinsically present in the SC neurons, and the later phase may use additional form information that could potentially be relayed to the SC from other brain areas (Fig. 2C, F).

We also analyzed microsaccades to remove potential eye movement confounds from our analyses. Microsaccade rate exhibited expected modulations as a function of time from image onset (Fig. S2A, E) [38-40]. This meant that in the early visual burst intervals of neural responses, there were already rare microsaccades due to microsaccadic inhibition. This ruled out a potential role for microsaccades in at least explaining the early visual burst interval results so far. However, we still repeated all analyses when excluding all trials containing microsaccades in the interval between -100 ms and +300 ms from image onset. Our results were largely unchanged (Fig. S2B-D, F-H). In fact, the AUC discrimination performance improved slightly across the board (compare Fig. S2B-D, F-H to Fig. 2), as might be expected given that microsaccades can modulate SC visual bursts [41, 42], and also given that these movements can cause measurable visual reafferent SC neural modulations after image jitter [43].

### Even visual-motor SC neurons detect objects in their very first visual responses

To further appreciate the SC’s role in express object detection, even within the initial visual bursts, we also considered this structure’s different functional neuron types. For example, it is well known that deeper-layer visual-motor neurons are relevant for a variety of cognitive processes like target selection, attention, and decision making [18-20, 22-24, 44], in addition to their roles in eye movement generation [45-48]. So, we functionally classified our neurons according to classic visual and saccade-related response criteria (Methods), and we then explored object detection performance once again.

In both monkeys, most of our neurons were visual-motor-prelude neurons (Methods): they emitted visual bursts after stimulus onset, saccade-related bursts at saccade onset, as well as significant prelude activity (above baseline spiking rate) before saccade onset. We also encountered visual-motor neurons, which did not exhibit substantial delay-period (prelude) activity but were otherwise similar to visual-motor-prelude neurons. Finally, our database included a fewer number of purely visual neurons, which came in two primary flavors: visual neurons emitting a burst shortly after stimulus onset, and visual-delay neurons also exhibiting delay-period activity after the bursts.

All neuron types that we encountered exhibited significant object detection capabilities, and highly similarly in both monkeys. For example, Fig. 3A, D shows the distribution of neuron types contributing to the results of Fig. 2A, D. Both visual-motor types were most frequent in both monkeys (likely due to the recording technique with thick electrode shanks; Methods), but purely visual neurons were also clearly present. Most interestingly, visual-motor neurons detected visual objects even earlier than visual-delay neurons in both monkeys (with the caveat that the number of the visual-delay neurons was relatively low). This result is illustrated in Fig. 3B, E: in both animals, visual-motor-prelude neurons exhibited high AUC discrimination performance (relative to grid-scrambled images) in their very initial visual bursts, and this high discrimination performance actually preceded the discrimination performance of visual-delay neurons. Even though the numbers of visual-delay neurons were relatively low in each animal, the effects in both animals were virtually identical, increasing our confidence in concluding that there is indeed very early object detection by visual-motor-prelude neurons. At the very least, it is safe to state that visual-motor-prelude neurons detect visual objects as early as (if not earlier) than purely visual neurons (Fig. 3B, E; also see Fig. 3C, F).

**Figure 3.**
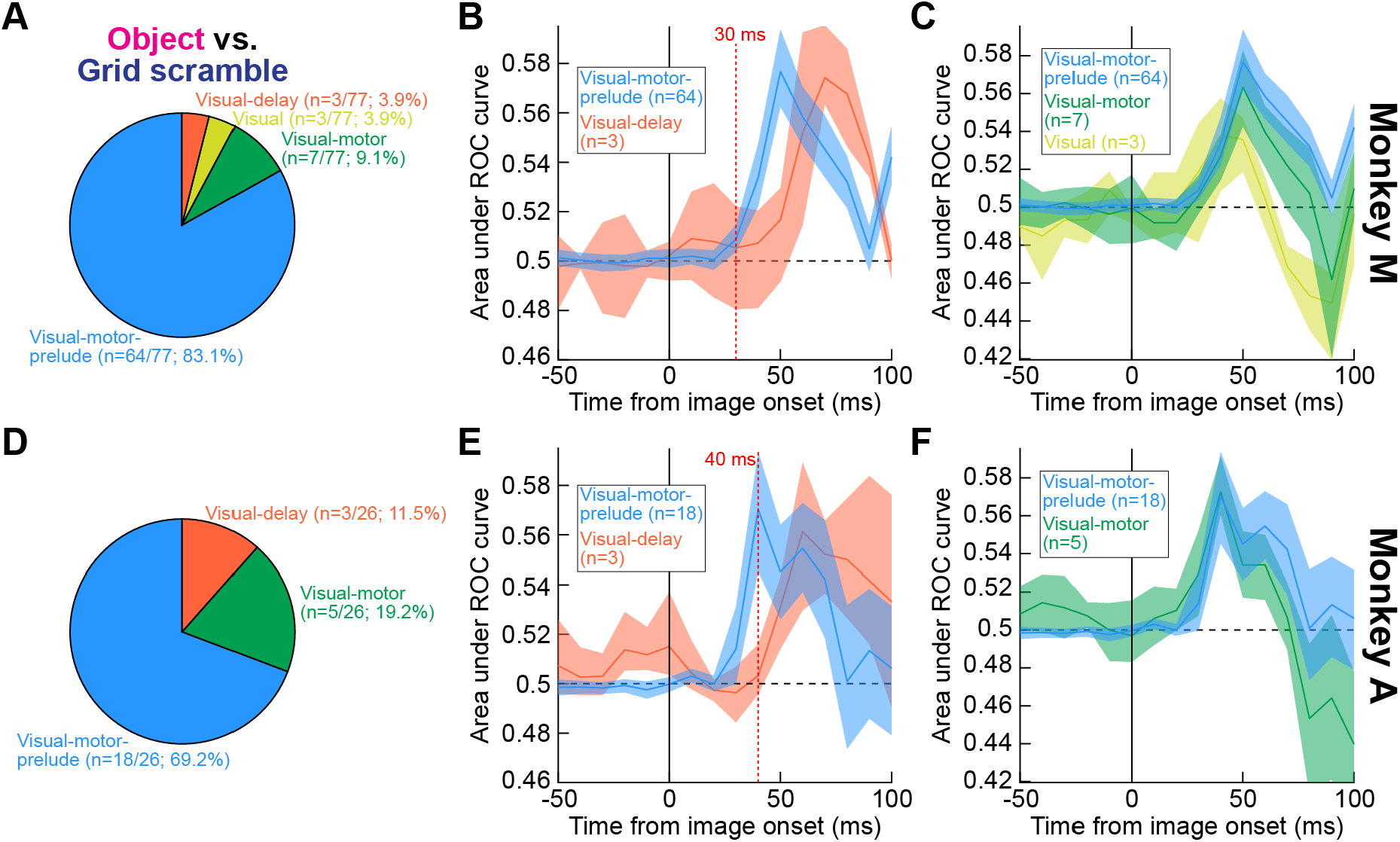
Express object detection even by visual-motor neurons. **(A)** Distribution of neuron types (Methods) exhibiting significant object detection performance in the data of Fig. 2A (monkey M). Visual-motor and purely visual neurons were both present. **(B)** When we compared the initial AUC discrimination performance between visual-motor-prelude and visual-delay neurons, we found earlier object detection by the visual-motor-prelude neurons (with the caveat of significantly fewer visual-delay neurons in the database). Error bars denote 95% confidence intervals. **(C)** Similarly, visual-motor neurons (without prelude activity) also exhibited early detection performance. In this panel, the curve from visual-motor-prelude neurons is replicated from **B** to facilitate comparison. In this animal, a few visual neurons were also encountered that exhibited object detection performance, and their results are shown in yellow. Thus, all visual and visual-motor neuron types detected objects in this animal, and it is interesting that even visual-motor neurons exhibited early detection. **(D-F)** Highly similar results from monkey A. Note that in this monkey, we did not encounter visual neurons, so they are not shown in **F** as they were shown in **C**. Figure S3 provides further analyses of neuron types, focusing on later, sustained intervals of neural discharge.

We also found that prelude activity was not a prerequisite for visual-motor neurons to exhibit rapid object detection. Specifically, in Fig. 3C, F, we repeated the ROC analyses but now for the visual-motor neurons (green), which did not have substantial delay-period activity. For comparison, we also plotted the visual-motor-prelude neuron results from Fig. 3B, E again, to facilitate comparing the curves. Both neuron types exhibited similar early detection of intact visual objects relative to grid-scrambled images (similar results were also obtained with phase scrambling). In monkey M, we also had some purely visual (burst) neurons, and they also exhibited early object detection (yellow in Fig. 3C). Therefore, all of the above results suggest that visual-motor SC neurons are a substantial contributor to the SC’s ability to rapidly detect visual objects.

In terms of later intervals after image onset, perhaps expectedly, the neurons that had sustained activity also showed sustained significant AUC discrimination performance between object and scramble images. For example, when we repeated the ROC analyses of Figs. 2, 3 for visual-motor-prelude and visual-delay neurons combined (both of which had sustained activity), and we compared them to visual-motor and visual neurons (both not having sustained activity), we found that the later (>100 ms) AUC discrimination performance was systematically higher for the former group of neurons (Fig. S3). This makes sense because sustained activity provides a necessary spiking substrate for encoding information about the visual images.

Therefore, not only do SC neurons detect visual objects early (Figs. 1, 2), but they do so even if they are motor-related neurons (Fig. 3). Moreover, delay-period activity contributes to maintaining information about the intact object images for sustained intervals, as might be the case in a variety of cognitive tasks.

### Individual SC neurons exhibit early and late preference for individual object categories

The results so far pooled all seven object categories presented to each neuron in the analyses (Methods). However, we also noticed that SC neurons can be differentially modulated by specific images. For example, inspection of the spike rasters of the neuron of Fig. 1B, which are grouped by object category, reveals that the neuron fired the most action potentials upon presentation of the neutral human face and the least action potentials after the neutral monkey face appeared. Therefore, not only did the neuron detect the presence of intact object images in its RF (Fig. 1B), but its response was also differentially modulated for different image categories. This motivated us to inspect visual object preference in more detail, and we did so using two approaches.

First, we took a strict approach of only analyzing neurons in which activity for any of the seven object categories (in the 0-300 ms interval after image onset) was significantly discriminable from all scrambled images (i.e. both the grid- and phase-scrambled images). If, and only if, a neuron satisfied this constraint, we defined the preferred object category as the category for which the peak AUC value in the interval 0-300 ms after image onset was higher than all other object categories. In monkey M, this resulted in 42 neurons (Fig. 4A), and in monkey A, we found 15 neurons satisfying these conditions (Fig. 4B). For each of these neurons, we plotted in Fig. 4A, B the AUC values for the preferred object relative to all scrambles. There was clear discriminability of the preferred objects from the control images. Most importantly, both monkeys exhibited elevated AUC values in the early visual burst intervals (much like in Figs. 2, 3 above). Therefore, object preference in the SC emerges quickly (within approximately 50 ms from image onset).

**Figure 4.**
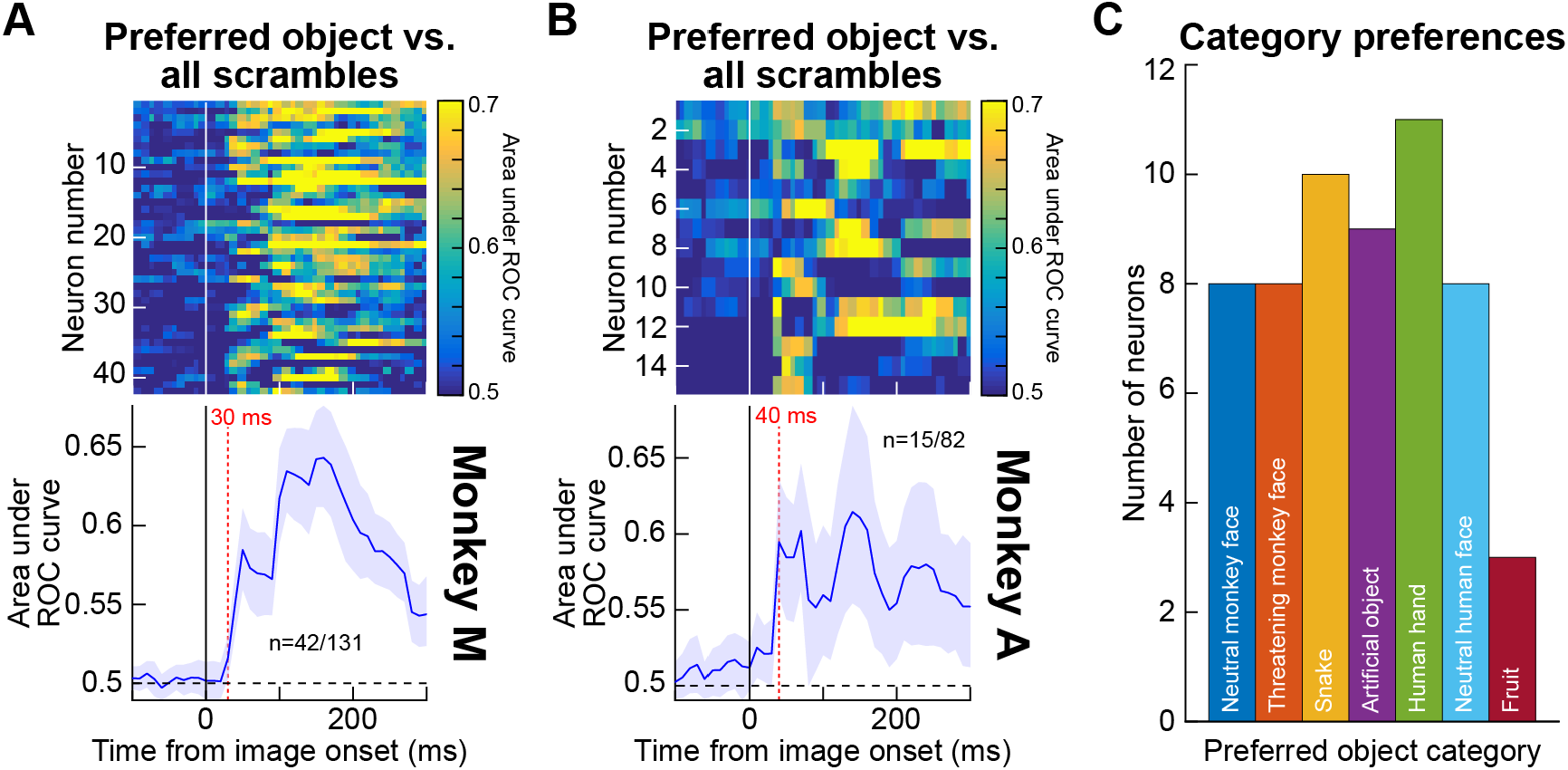
Early preference for specific object categories, and with a diversity of category preferences across the population. **(A)** ROC analyses in monkey M comparing firing rates for the most preferred object category to firing rates in both the grid- and phase-scrambled images (Methods). Object preference emerged even in the early visual burst interval of neural responses (<100 ms). All conventions are as in Fig. 2. **(B)** Similar results for monkey A. **(C)** Distribution of most preferred object in the analyses of **A, B**. There was a diversity of preferences across categories (individual monkey results are shown in Fig. S4).

Using the same approach, we also checked whether specific object categories were more or less frequently preferred. For example, it could be that threatening stimuli (e.g. snakes and threatening monkey faces) would be particularly relevant for object detection by the SC [12, 14]. On the other hand, a role for the SC in modulating cortical visual areas related to object recognition [29] might suggest the need for more diversity in the SC representation. Therefore, for each neuron in Fig. 4A, B, we checked which object category was actually preferred by the neuron (as per the same definition of object preference as in Fig. 4A, B). As shown in Fig. 4C, the distribution was diverse and without particular predominance of threatening objects (Fig. S4 shows individual monkey results). This implies a more generalized role of the SC in rapid object detection and discrimination than simply the flagging of threatening stimuli or of faces.

Our second approach to establish the presence of express and late object preference in SC neural discharge, as a general property, was to demonstrate a clear differential in firing rates for different objects, which disappeared when the objects were scrambled. For each neuron in the entire database, we picked the object category that resulted in the most or least visually-evoked action potentials; the object category was then labeled as the preferred or non-preferred object category accordingly. We also did this for either the early visual burst interval (0-100 ms from image onset) or the sustained interval (100-300 ms). By definition of the analysis, there was a robust firing rate difference between the preferred and non-preferred object categories. We then took the same categories and compared the firing rates in the grid-scrambled versions of the same images. If the difference in firing rate between preferred and non-preferred objects was due to low-level image features, then this difference should have persisted even in the grid-scrambled image comparisons. If not, it would suggest that there were indeed preferred and non-preferred visual form images in the individual session for the neurons.

SC neurons demonstrated preference for specific object images even in their very initial visual bursts. Consider, for example, the neuron shown in Fig. 5A, which is the same as that in Fig. 1F. In the left column, we plotted the neuron’s responses to the preferred (neutral human face) and non-preferred (neutral monkey face) object images. As per the definition of the preferred and non-preferred analysis, there was a clear difference in initial visual burst strength (the shaded gray region shows our “early” analysis interval). Most critically, this difference disappeared for the grid-scrambled versions of the images (right column), and the visual burst strength for the grid-scrambled images was lower than the firing rate for the preferred object image (consistent with Figs. 1-4). Therefore, something about the intact object images, which was not present in the scrambles, was relevant for the response of the neuron to differentiate between the human (preferred) and monkey (non-preferred) faces.

**Figure 5.**
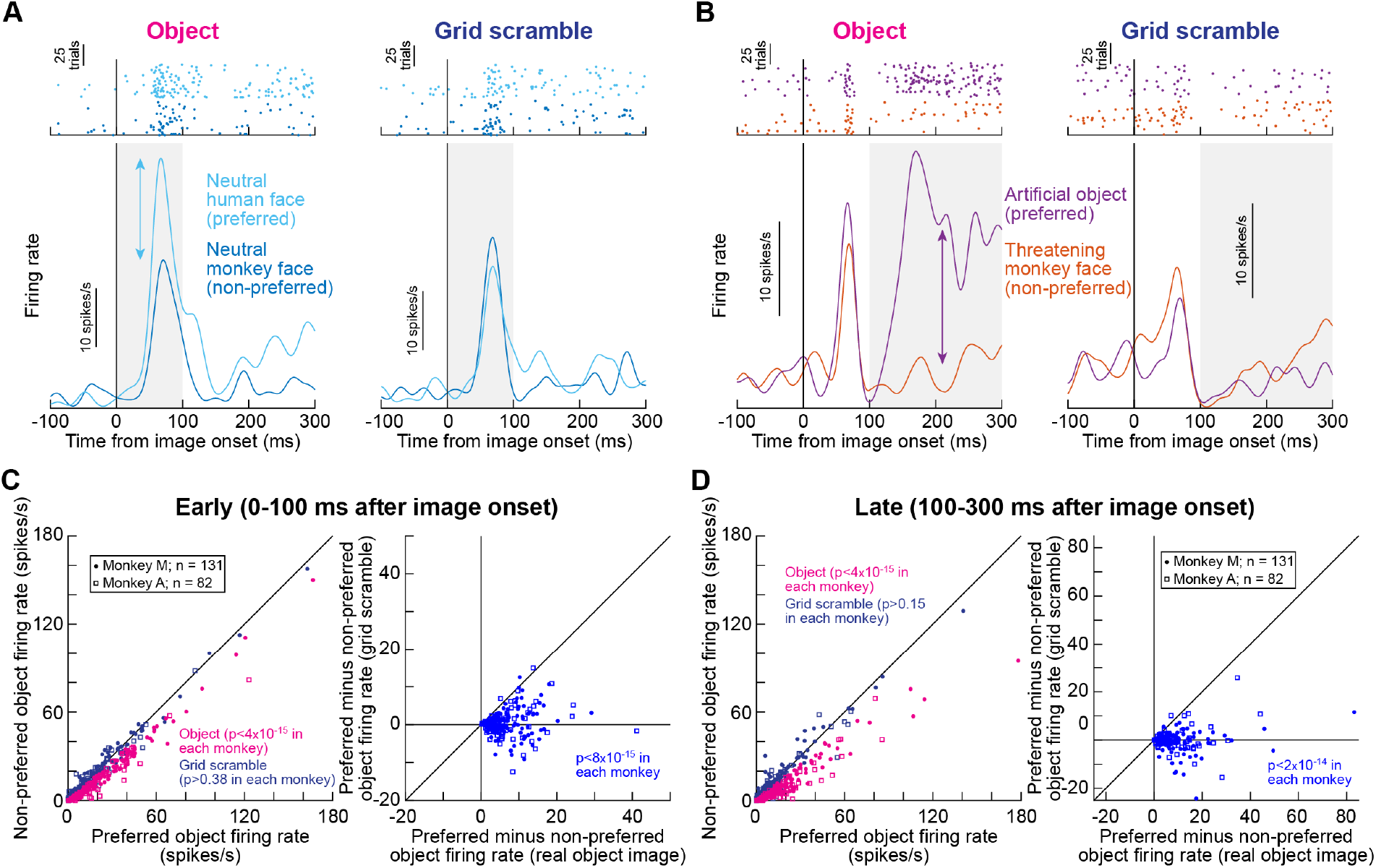
Discrimination of object categories by SC neurons even in the very initial visual responses. **(A)** Responses of the neuron shown in Fig. 1F for the most and least preferred object categories in the session (left column). In the right column, the difference in visual burst strength evident for the intact object images (left column) disappeared. The gray shaded region marks our analysis interval for assessing object preference in the early visual burst intervals. **(B)** A second neuron (same as that in Fig. 1C) exhibiting object preference in the sustained interval (shaded region), which again disappeared under grid scrambling (compare left and right columns). Note that the two neurons preferred different object categories, consistent with Fig. 4C. **(C)** Summary across all isolated neurons in our database of preferred and non-preferred early responses, with both intact and grid-scrambled images (color coded in the left panel). The left panel shows raw measurements, and the right panel plots differences of firing rates between preferred and non-preferred conditions (intact images on the x-axis and scrambled images on the y-axis). In both cases, we used a Wilcoxon signed rank test within each monkey for statistical testing. Real object tuning (significant differences in the left panel and >0 x-axis values in the right panel) disappeared when the same images were grid-scrambled. **(D)** Same as **C** but for the late, sustained interval, and with similar conclusions.

This second analysis approach avoided the caveat in Fig. 4A, B that the preferred object could have only had the peak AUC value much later than in the initial visual burst. However, we also checked for object preference with this second approach in the sustained firing rates of the neurons as well. For example, the neuron in Fig. 5B is the same as that in Fig. 1C, but we now inspected individual object categories. This neuron clearly preferred the artificial object image in its sustained visual response (shaded gray region delineating our “late” analysis interval), and its least preferred image for the session was the threatening monkey face (left column). Again, most critically, this difference disappeared when comparing the grid-scrambled versions of the same images (right column). Therefore, both early and late intervals demonstrated object preference by SC neurons.

Across the population, a clear difference in responses to preferred and non-preferred object images within a given session was absent with grid-scrambling, and this happened even in the early visual burst interval (Fig. 5C). The left column in Fig. 5C shows preferred and non-preferred responses for the intact object images and for the grid-scrambled images. There was a significant difference (Wilcoxon signed rank test; p<4×10^−15^) only for the intact object images, as also clarified in the right column plotting the difference response between preferred and non-preferred objects under the two conditions (each monkey’s results are shown individually with different symbols). A similar result was also seen for the late sustained interval (Fig. 5D; Wilcoxon signed rank test; p<4×10^−15^ for the real images and p>0.15 for grid scrambling). Therefore, in all, our results demonstrate robust, express detection (Figs. 1-3) and discrimination (Figs. 4, 5) of visual objects by primate SC neurons.

## Discussion

We found that all of our classified visual and visual-motor SC neuron types contributed to rapid detection and discrimination of visual objects, with even deeper visual-motor neurons doing so in their very first visual bursts. Such visual-motor neurons are typically implicated in a variety of cognitive functions beyond saccade generation [18-20, 49-51], suggesting that the visual form information that they carry can influence such functions as well. Moreover, because of the intrinsic motor nature of these neurons, it would also be very intriguing to think of the role of the SC’s visual object representations in the broader context of active vision with saccades.

Besides the short latencies associated with object detection and discrimination by SC neurons, we were particularly intrigued by the role of spatial frequency information in object detection during the early phases of SC neural responses. In the early visual burst phase of neural responses, we observed that AUC discriminability between the responses for object images and spectral-matched phase-scrambled images was weak (Fig. 2C, F). This suggests that a functional role for spatial frequency tuning in SC neurons [31] could be to aid in rapid object detection. Indeed, this could also mediate rapid orienting responses to objects [8], since the spatial frequency tuning of SC neurons is relevant for saccadic reaction times [31]. Having said that, spatial frequency information cannot fully explain early object detection by the SC because the AUC discrimination performance between intact objects and phase-scrambled controls still became significant earlier than 100 ms after image onset (Fig. 2). This is still relatively faster than when some cortical visual areas detect objects [52], again affirming a role for the SC in early object detection. This is also consistent with early pop out of high level visual objects, like faces, in perception [7].

Another interesting observation is that object detection in later intervals after the visual bursts (e.g. >100 ms after image onset in the phase scrambling results) seems to rely on more than just the spatial frequency information. This is because AUC discrimination was still significant between object and phase-scrambled controls in these later intervals (Fig. 2C, F), and it would imply potential feedback from other visual cortical areas involved in object processing. This could functionally allow visual cortical areas to utilize additional spatial frequency bands, and other rich visual feature representations, beyond those represented in the SC. That is, since SC neurons are predominantly low-pass in nature at our tested eccentricities [31] (Fig. S5), and since various cortical areas can detect objects at multiple spatial frequency bands [33], feedback from these areas could help to sustain the object representations in the SC after the initial visual bursts subside. This is important because object recognition does indeed benefit from middle spatial frequencies in images [34, 35, 37].

We are also intrigued by the object preference results, especially in the earliest phases of neural responses (Figs. 4, 5). In previous work, it was suggested that the SC is part of a network for quickly detecting threats and/or faces [6]. Indeed, SC lesions in infant monkeys impair these monkeys’ fear responses to snakes [12]. However, the SC seems to influence cortical visual processing in a more generalized manner [8, 29, 30], suggesting that there is value in having the SC act as a more generalized object detector and discriminator as opposed to only a face and threat detector. This is consistent with our observations; we found a diversity of preferred object images across the population. Of course, very fine discriminations may be ultimately limited by the potential pattern processing capacity limitations of SC neurons, such as orientation [53] and spatial frequency [31] bandwidths, but some level of “recognition” by the SC may still be useful for facilitating orienting responses to objects in our environment.

In all, our results motivate further investigations of subcortical pathways for visual perception, particularly given the active nature of behavior in the real world, and the perpetual interplay between sensory processing, on the one hand, and movement generation, on the other.

## Acknowledgements

We were funded by the Deutsche Forschungsgemeinschaft (DFG) through projects: (1) BO5681/1-1; and (2) SFB 1233, Robust Vision: Inference Principles and Neural Mechanisms, TP 11, project number: 276693517.

## Author contributions

Both authors conceived the study. ARB collected the data. Both authors interpreted the results and wrote the manuscript.

## Methods

### Experimental animals and ethics approvals

We recorded superior colliculus (SC) neural activity from two adult, male rhesus macaque monkeys (A and M) aged 9 and 8 years, respectively. The experiments were approved by ethics committees at the regional governmental offices of the city of Tübingen.

### Laboratory setup and animal preparation

The experiments were conducted in the same laboratory as that described for the monkey portions of [8]. Briefly, the monkeys were seated in a darkened booth ∼72 cm from a calibrated and linearized CRT display spanning ∼31 deg horizontally and ∼23 deg vertically. Data acquisition and stimulus control were managed by a modified version of PLDAPS [54], interfacing with the Psychophysics Toolbox [55-57] and an OmniPlex data acquisition system (Plexon, Inc.).

The monkeys were prepared for behavioral training and electrophysiological recordings earlier [58, 59]. Specifically, each monkey was implanted with a head-holder and scleral search coil in one eye [58]. The search coil allowed tracking eye movements using the magnetic induction technique [60, 61], and the head-holder comfortably stabilized head position during the experiments. The monkeys also each had a recording chamber centered on the midline and tilted 38 deg posterior of vertical, allowing access to both the right and left SC.

### Behavioral task

We employed a modified version of the classic delayed, visually-guided saccade task, similar to what we did in our recent behavioral study [8] (see Fig. S1). Each trial started with the appearance of a central white fixation spot of 79.9 cd/m^2^ luminance, presented over a gray background (26.11 cd/m^2^). The fixation spot was 0.18 × 0.18 deg in dimensions. After 300 ms, an image patch (see below for image preparation procedures) appeared within the visual response fields (RF’s) of the recorded neurons. The image patch could contain pictures of real-life objects or the other versions of image controls described in more detail below. After 300-700 ms from image patch onset, a white spot identical to the fixation spot appeared on top of a gray disc (diameter: 0.54 deg; 26.11 cd/m^2^) in the center of the image patch. This white spot was referred to as the saccade target in our analyses. It remained visible (along with the fixation spot and image patch) for 500-1000 ms, at which point the fixation spot disappeared to instruct the monkeys to generate a saccade towards the saccade target (and the underlying image patch). If the monkey successfully made the saccade within 500 ms, it received positive reinforcement in the form of liquid reward.

As described in more detail below, the size of the image patch that we presented was matched to the RF size, and its position was designated after initial assessment of RF locations and sizes (using standard visual and saccadic tasks employed in SC studies; our instantiations of these tasks were described previously [59, 62]). The average luminance of the image patch was 42.07 cd/m^2^.

### Image database and image pre-processing procedures

We used a total of 156 grayscale images, from previously published studies [8, 29, 63], across seven different object categories: neutral monkey face (15 images), threatening monkey face (15 images), snake (15 images), artificial object (15 images), human hand (16 images), neutral human face (64 images), and fruit (16 images) (Figs. 1A, S1). In each session, we randomly picked seven images from the database, one from each category.

For each session, we first sized the images to match the RF sizes of the neurons across the recording contacts. Our neurons spanned eccentricities in the range of 3.1-23.9 deg (Fig. S5), and we assessed their RF’s using standard visual and saccadic tasks. The image patches were square, and their sizes were in the range of 2-8 deg (in width and height). These sizes fit within the excitatory parts of the neurons’ RF’s. Since we had multiple RF’s within a session (see neurophysiological procedures below), we picked the image location that best matched most of these RF’s. This was feasible given the topographic organization of the SC and the fact that our electrode penetrations were roughly orthogonal to the SC surface at our recorded eccentricities.

We then iteratively equalized the luminance histograms and spatial frequency spectra of the seven images of a given session using the SHINE toolbox [64]. Specifically, we ran 20 iterations of histogram matching (*histMatch* function) of the gray levels across the images, as well as spectral matching across the same images (*specMatch* function). To generate phase-scrambled images, we randomized the phase matrices of the Fourier-decomposed images, while keeping the amplitude matrices unchanged. Then, to match the real and phase-scrambled images further, we took all object images and their corresponding phase-scrambled images, and we again iteratively matched them once more for histogram levels and frequency spectra using the same SHINE toolbox functions (again, with 20 iterations). Example final images (real and phase-scrambled) are shown in Fig. 1A and Fig. S1B.

To obtain the grid-scrambled image controls, we overlaid 1-pixel-width horizontal and vertical lines of mean image luminance over the real object images. These horizontal and vertical lines formed a grid of 0.33 deg x 0.33 deg squares within which the original object was visible. We then scrambled all grids by randomizing their original locations in the image. To ensure that the neural modulations associated with the grid-scrambled images were not fully explained by the overlaid horizontal and vertical gray lines, we also created the grid overlay without randomizing the individual grid locations. This created the object+grid images (as if the objects were intact and only occluded by a thin grid in front of them). Examples of the final grid-scrambled and object+grid images used in our study are shown in Fig. 1A and Fig. S1B.

### Neurophysiological procedures and functional cell type classification

We recorded neural activity using linear microelectrode arrays (V-Probes, Plexon, Inc.) inserted into the SC. We aligned the arrays (16- or 24-channels with 50 μm inter-electrode spacing) to obtain sufficient coverage across different functional SC layers (0.8-1.2 mm depth coverage by the contacts).

The experiment started by identifying entry into the SC by the deepest electrode contact, and we then advanced the array to insert further contacts into the SC. After ensuring that the tissue had settled and the neural activity was stabilized across contacts, we assessed the RF’s at the electrode contacts using standard visual and saccade tasks. This allowed us to place and size the object images for a given session according to the neurons’ approximate RF locations and sizes. Following RF estimation and the preparation of the object and control images to fit the RF sizes, we ran the main experiment and collected an average of 32 (+/-8 SD) trial repetitions per session of the different image conditions that we had: 4 image patch versions (real object, phase-scrambled, grid-scrambled, and object+grid) of each of the 7 object categories (total of 28 different conditions), resulting in a total of 903 (+/-239 SD) trials per session.

We classified neurons as being visual, delay, visual-delay, visual-motor, visual-motor-prelude, or motor in nature, as per previous criteria [65]. Specifically, in our delayed visually-guided saccade to image task, we measured the firing rate in each trial, regardless of image conditions, during four different epochs: baseline (100 ms before image onset), visual (50-150 ms after image onset), delay (400-500 ms after saccade target onset), and motor (−50 to 25 ms from saccade onset). Next, we used the firing rates in these four epochs to compute a non-parametric ANOVA (Kruskal-Wallis), and we determined the neuron class by post-hoc significance tests (p<0.05). Neurons with significant activity in the visual epoch compared to the baseline epoch were classified as visual neurons. Similarly, neurons with significant activity in the motor epoch compared to the baseline and delay epochs were classified as motor neurons, and a visual neuron with significant motor activity was classified as a visual-motor neuron. Furthermore, visual neurons posessing significant delay-period activity were labeled as visual-delay neurons, and visual-motor neurons with significant delay-period activity were classified as visual-motor-prelude neurons. Any neuron that did not have higher than 5 spikes/s average firing rate in any of the above-mentioned measurement intervals (other than baseline) was excluded from further study. Similarly, for the purposes of this study, we did not analyze the purely motor neurons, since we were interested in assessing visual object detection by the SC.

In total, we had 82 included neurons from monkey A and 131 from monkey M. Approximately half of the neurons in monkey A (47.56%) and two thirds in monkey M (67.18%) were visual-motor-prelude neurons in our database. The next most frequent neuron type in our sample was visual-motor neurons (19.51% in monkey A and 16.03% in monkey M), followed by the motor (13.41% and 6.87%) and visual-delay (13.41% and 5.34%) neurons, and then finally the visual neurons (6.1% and 3.82%). Delay-only neurons were a rarity (1 in monkey M and non-existent in monkey A), and were not analyzed. The neurons’ preferred RF hotspot locations are shown in Fig. S5.

### Data analysis

We detected saccades and microsaccades using our previously described toolbox [66], and we inspected the detection results manually. To investigate whether microsaccades at image onset might have influenced the SC responses to the stimuli, whether by peri-microsaccadic modulation [41, 62] or jittering of images [43], we computed microsaccade rate across time from image onset (e.g. Fig. S2A, E). We did so similarly to how we estimated microsaccade rate recently [8]. Briefly, we binned microsaccades using a 40 ms moving time window, with time steps of 10 ms. In general, we included all trials in our neural data analyses, even when there were microsaccades. This was fine because of the low likelihood of microsaccades, especially in the critical early visual burst interval. However, we also confirmed that our results were unchanged by repeating the analyses after removing all trials in which there was a microsaccade between -100 ms and +300 ms relative to image onset (e.g. Fig. S2).

For neural analyses, we sorted the neurons offline using the Kilosort Toolbox [67], followed by manual curation using the phy software. We then proceeded to analyze the spike rasters and firing rates.

To investigate whether SC visual responses differentiate between object and non-object stimuli, we plotted spike rasters and firing rates across the different image conditions (e.g. Fig. 1). We then assessed whether an ideal observer could discriminate between object and non-object stimuli just based on the SC firing rates. To do so, we performed receiver operating characteristic (ROC) analyses using 40 ms time bins moving in steps of 10 ms. In each 40 ms time bin around the time of image onset, we collected firing rates within this interval from all trials of the real-life object condition and all trials of an image control from the same neuron (e.g. phase-scrambled or grid-scrambled images). We then ran the ROC analysis to obtain an area under ROC curve measure (AUC), allowing us to assess the discriminability between the two firing rate distributions. An area under the ROC curve value of 0.5 would indicate non-discriminable firing rate distributions. We performed the ROC analyses at all times from -100 ms to +300 ms from image onset, with 10 ms resolution. We did this because the earliest time at which the saccade target could appear in the task was 300 ms (e.g. Fig. S1). We assessed a neuron as detecting objects if its area under the ROC curve in any interval between 0 and 300 ms was statistically significantly different from 0.5. We assessed significance by calculating bootstrapped confidence intervals for the area under the ROC curve measure and using a p<0.05 criterion. This is similar to our previous approaches [29]. We then averaged across all significant neurons’ AUC time courses and obtained 95% confidence intervals across the population. We labeled the time of object detection in figures as the time at which the population AUC discrimination time course first deviated significantly from 0.5 (i.e. no overlap between the 95% confidence interval and 0.5).

We also repeated the ROC analyses for the different functionally-classified neurons. For example, we picked only visual-motor-prelude neurons and calculated the area under the ROC curve metrics for those, or we only considered visual-delay neurons. This allowed us to assess whether early visual object detection by the SC (e.g. in the initial visual burst interval; see Results) only occurred in purely sensory neurons, or whether it also appeared in deeper visual-motor neurons. In some analyses, we found that whether a neuron had delay-period activity or not (e.g. visual-delay and visual-motor-prelude neurons both had delay-period activity) influenced the ROC results in either early or late intervals after image onset. Therefore, to demonstrate this point, we combined neuron types appropriately; that is, visual-motor-prelude and visual-delay neurons were combined together since they both showed delay-period activity, and visual-motor or visual neurons were combined together because they both lacked delay-period activity.

To assess whether SC neurons could also discriminate between different object categories presented within a given session, we investigated object preference in a variety of ways. For each neuron, we first plotted firing rates as a function of object category (example neurons are shown in Figs. 1, 5). We found that different neurons have higher firing rates for different objects, whether in the initial visual burst interval or in the later sustained response (e.g. Fig. 5). To analyze such preference further, we first looked at the strict criterion of only those neurons exhibiting significant AUC results for individual object images with respect to all scrambled images. Therefore, for each neuron, we performed ROC analyses comparing responses to individual object images with responses to all scrambled images (i.e. both phase- and grid-scrambled conditions). The preferred object of a given neuron was labeled as the object with the significant and highest AUC value in any time interval 0-300 ms after image onset. Across the population, we then checked whether specific object categories were more or less prevalent as the preferred objects of the neurons.

The above approach allowed us to look at object preference using a strict measure that captures the difference in activity between real object images and control scramble images. This way, we could simultaneously conclude that (1) the neurons detected objects as opposed to non-object control images, and (2) the same neurons exhibited a preference for certain objects as opposed to others. However, the peak AUC value could appear anywhere in the first 300 ms, and we were particularly interested in whether there was object preference in the very earliest visual bursts. Therefore, we also checked for the existence of object preference by SC neurons using another approach. For all of the original categorized neurons in each animal, we picked either an early visual burst interval (0-100 ms from image onset) or a late sustained interval (100-300 ms from image onset). In each interval, we picked, for a given neuron, the object category that elicited the highest average firing rate (e.g. neutral human face). This was labeled the preferred object of this neuron. We also picked the object category evoking the lowest average activity in the same interval (e.g. neutral monkey face), and we labeled it as the non-preferred object. We then checked whether the difference in firing rate between the most and least preferred objects disappeared (or was significantly reduced) when the object images were scrambled. If the neurons were tuned to specific object categories, then the firing rate differences between preferred and non-preferred object images were expected to be higher than the differences in firing rates between the scrambled versions of these same objects. We statistically assessed such differences across the population using signed rank tests.

In all figures and analyses, we showed results for each monkey individually.

## Supplementary figures

**Figure S1.**
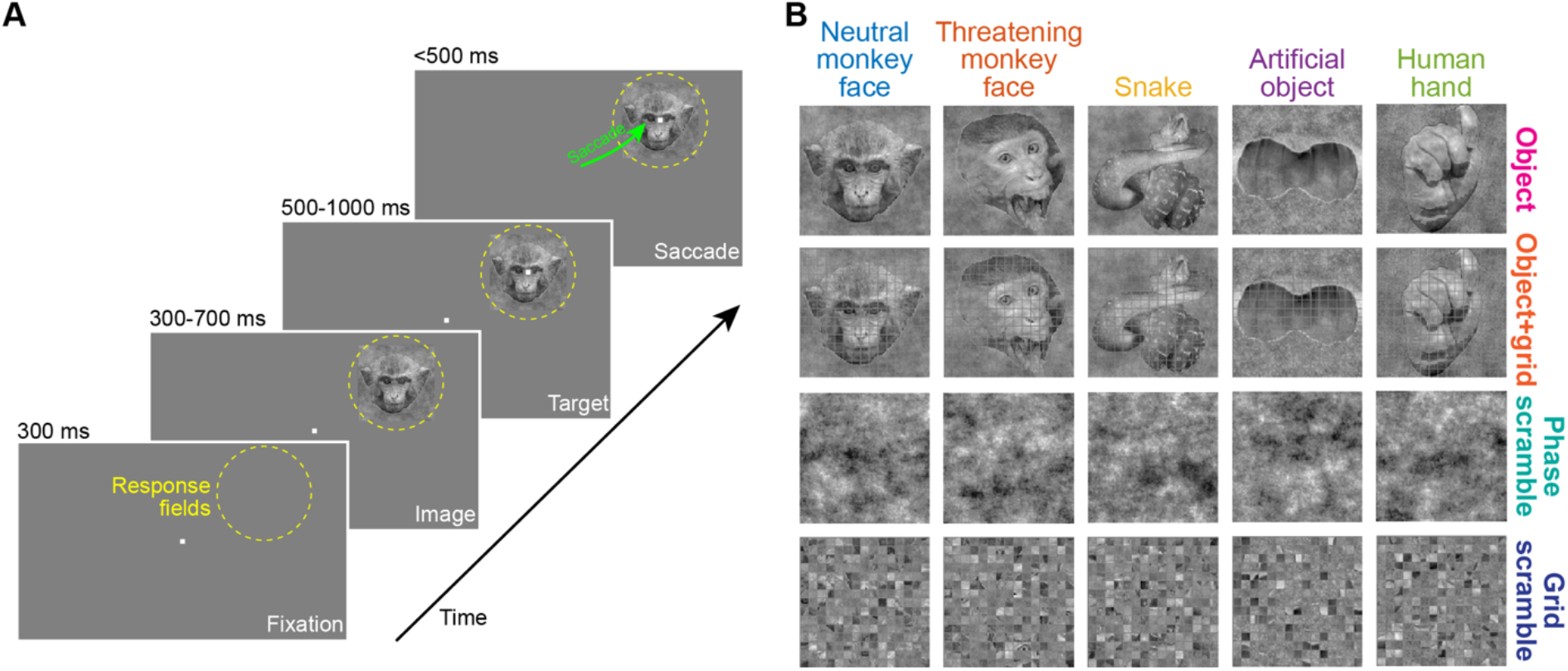
Behavioral task and example images. **(A)** Each task started with a central fixation spot. At the approximate response field (RF) locations of the recorded neurons in a given session (yellow dashed circle), an image appeared during fixation. After 300-700 ms from image onset, a saccade target appeared on top of the image for another fixation interval (500-1000 ms). The fixation spot then disappeared, instructing the monkey to generate a saccade towards the target on top of the image (green arrow). **(B)** Example images from a given session. The fruit image from the session is shown in Fig. 1, and the human neutral face image is not shown for data privacy reasons. The top row shows the real object images, and the second row shows these images with the grid overlay. The third row shows the phase-scrambled images, and the bottom row shows the grid-scrambled images (Methods).

**Figure S2.**
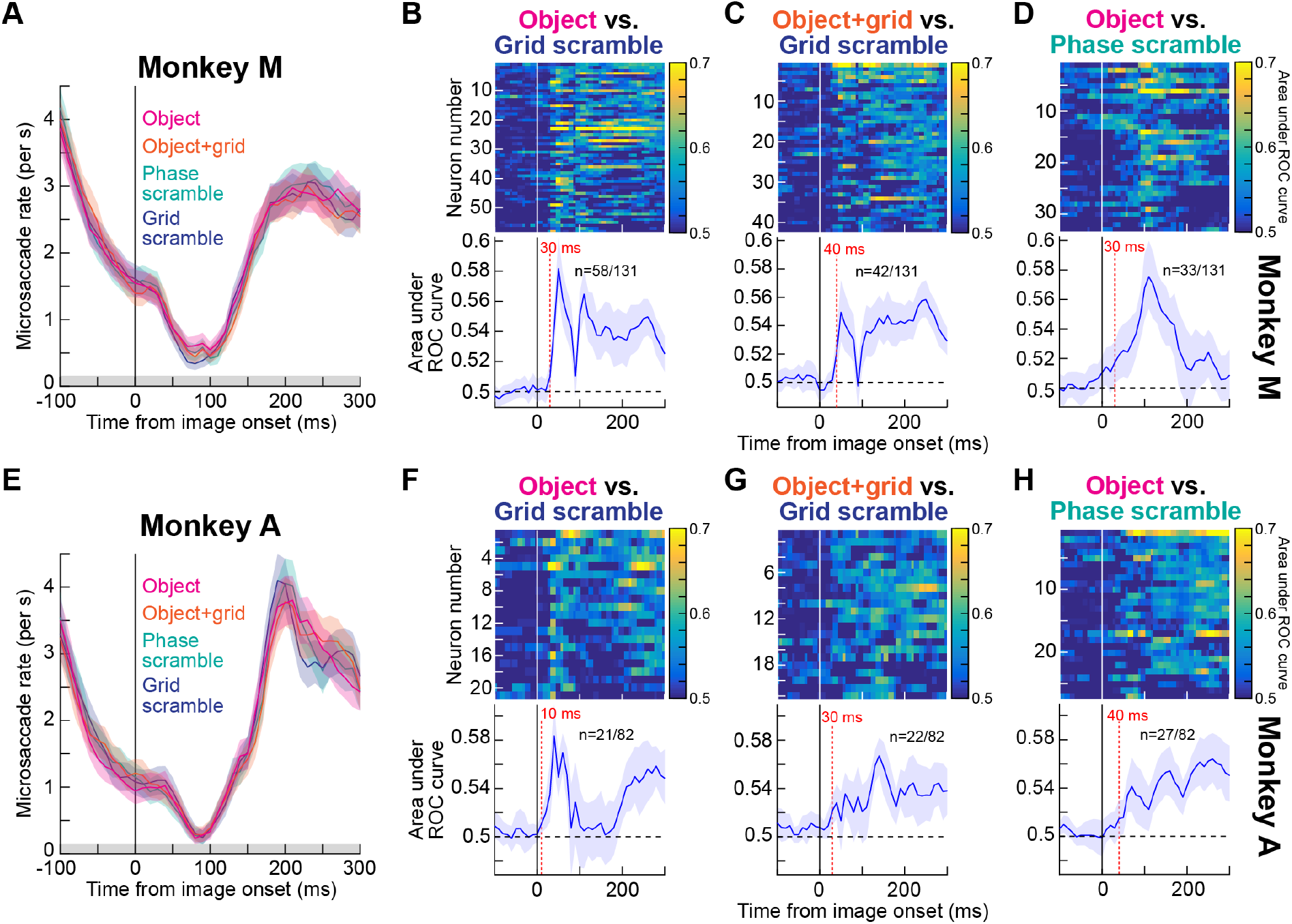
Discrimination between object and non-object stimuli by SC neurons’ visual responses, after controlling for microsaccades. **(A)** Microsaccade rate around the time of image onset from monkey M. A classic modulation of eye movement rate was present [38, 39, 68, 69]. Note that microsaccade rate was negligible in the early visual burst interval of neural responses, due to the known phenomenon of microsaccadic inhibition. The relatively high (but declining) microsaccade rate before image onset was due to the short initial fixation interval of the task (Fig. S1), and therefore had some refixation saccades as the monkey was starting a new trial after the end of the previous one. The gray bar on the x-axis denotes the interval chosen for removing microsaccades in the control analyses of **B**-**D. (B-D)** Same results as in Fig. 2A-C but after including only trials in which there were no microsaccades in the entire shown interval in **A** (−100 ms to +300 ms from image onset). The same qualitative results as in the main text were obtained. In fact, the AUC values here were generally higher than with all trials included. This is expected because microsaccades jitter images, and are associated with various effects on SC neurons’ firing rates [41-43, 62]. **(E)** Same as **A** but for monkey A. **(F-H)** Same as **B**-**D** but for monkey A.

**Figure S3.**
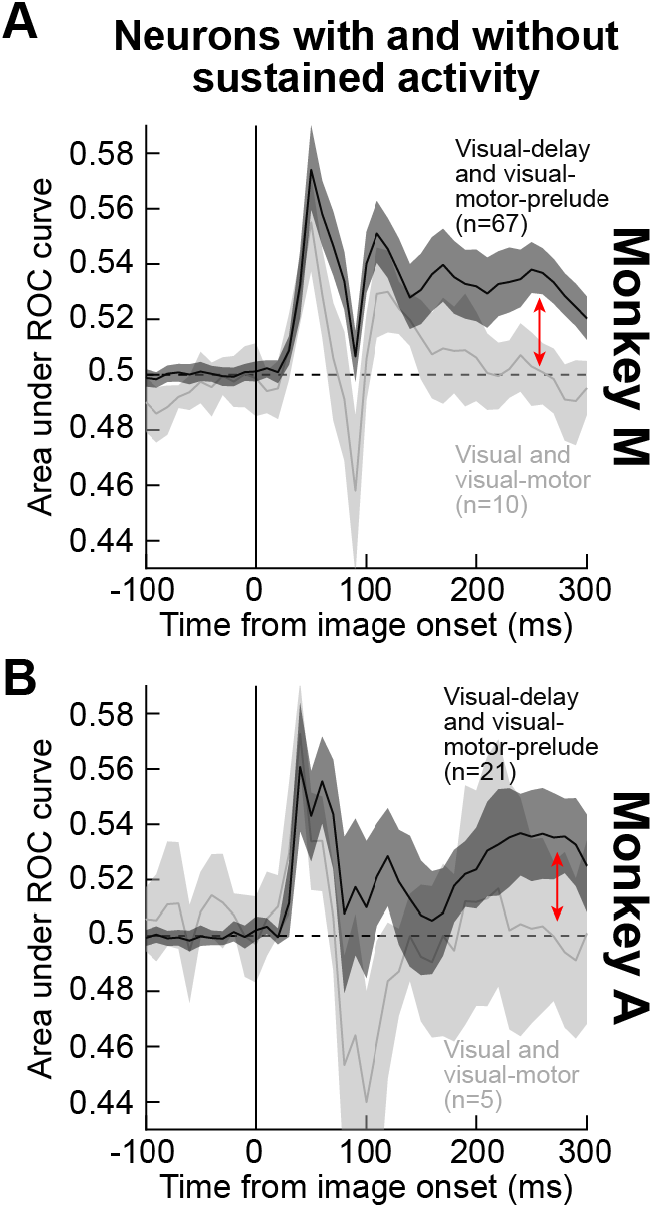
Neurons with sustained delay-period (prelude) activity allow sustained discrimination between object and non-object images in SC RF’s. **(A)** We performed our ROC analyses on object versus grid-scrambled images as in Figs. 2, 3, but this time by pooling only neurons with delay-period activity (visual-delay and visual-motor-prelude neurons) or neurons without (visual and visual-motor neurons). In the latter group, discrimination performance returned to baseline (light gray), whereas it remained significant throughout the sustained interval for the first group of neurons (see red vertical arrow). Error bars denote 95% confidence interval. **(B)** We observed very similar results in monkey A, although the smaller number of visual and visual-motor neurons (light gray) reduces the statistical confidence around this latter group of neurons.

**Figure S4.**
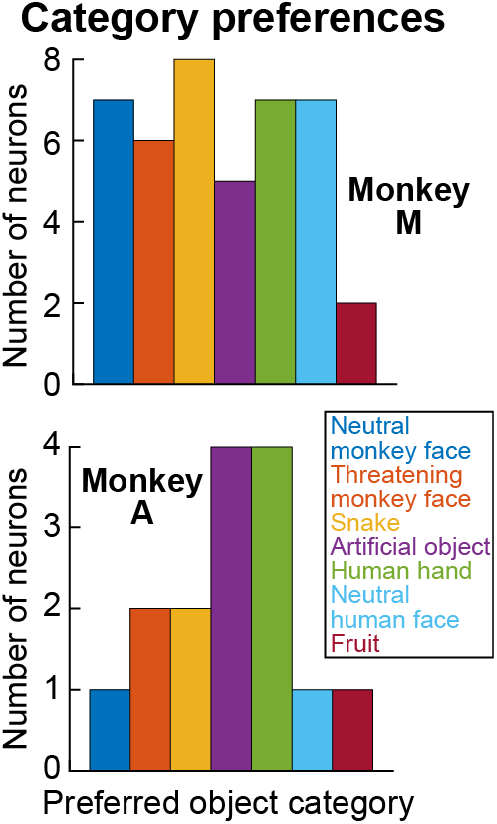
Individual monkey results from Fig. 4C. Each histogram shows the number of neurons preferring a given object category from each monkey, from the same analyses of Fig. 4. These neurons were, therefore, only the neurons that passed the AUC criterion relative to scrambled images (see Methods and Fig. 4). In both monkeys, no single category (e.g. snake or threatening monkey face) emerged as an outlier. Rather, there was diversity of object preferences, consistent with the idea of supporting object detection in general.

**Figure S5.**
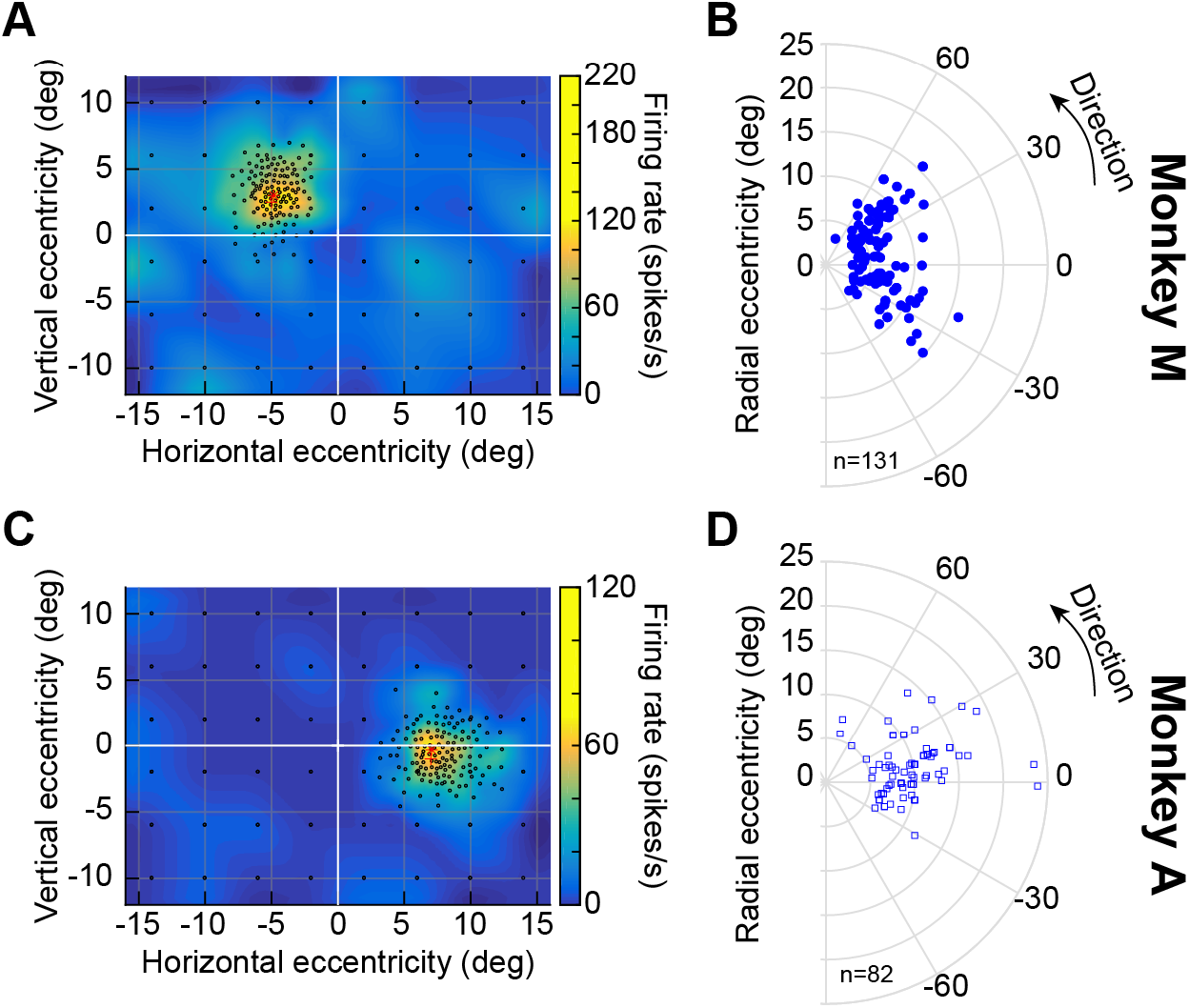
Response field (RF) locations of the recorded neurons. **(A)** Visual RF of an example neuron recorded from monkey M. Each black circle indicates a sampled location in which we presented a small spot during fixation. The pseudocolor surface indicates the mean firing rate emitted by the neuron in a visual epoch 40-140 ms after spot onset (we interpolated across space between the sampled locations to obtain the pseudocolor surface). The neuron’s RF occupied the upper left quadrant, and our online estimate of its hotspot is indicated by the red asterisk. The red cross indicates where we placed the image during the main experiment. **(B)** All RF hotspot locations from monkey M (remapped to one hemifield for easier viewing). Our neurons were extrafoveal. **(C)** Visual RF of an example neuron recorded from monkey A. The same conventions as in **A** apply. The neuron occupied the lower right quadrant. **(D)** All RF hotspot locations from monkey A, showing similar coverage to monkey M. For purely motor neurons, RF hotspot locations in **B, D** were obtained from the saccade-related, rather than visual, responses.

